# Sleep Benefits Memory for Semantic Category Structure While Preserving Exemplar-Specific Information

**DOI:** 10.1101/143172

**Authors:** Anna C. Schapiro, Elizabeth A. McDevitt, Lang Chen, Kenneth A. Norman, Sara C. Mednick, Timothy T. Rogers

## Abstract

Semantic memory encompasses knowledge about both the properties that typify concepts (e.g. robins, like all birds, have wings) as well as the properties that individuate conceptually related items (e.g. robins, in particular, have red breasts). We investigate the impact of sleep on new semantic learning using a property inference task in which both kinds of information are initially acquired equally well. Participants learned about three categories of novel objects possessing some properties that were shared among category exemplars and others that were unique to an exemplar, with exposure frequency varying across categories. In Experiment 1, memory for shared properties improved and memory for unique properties was preserved across a night of sleep, while memory for both feature types declined over a day awake. In Experiment 2, memory for shared properties improved across a nap, but only for the lower-frequency category, suggesting a prioritization of weakly learned information early in a sleep period. The increase was significantly correlated with amount of REM, but was also observed in participants who did not enter REM, suggesting involvement of both sleep stages. The results provide the first evidence that sleep improves memory for the shared structure of object categories, while simultaneously preserving object-unique information.

## Introduction

Semantic knowledge allows us to infer unobserved properties of newly encountered objects and events, requiring a mastery of both the coherent properties shared among conceptually related items and the individuating properties that distinguish them [1, 2]. For instance, the concept *bird* arises when children learn that certain property sets — having wings, beaks, feathers, hollow bones, and the name “bird” — all co-occur together or not at all [2–4]. That is, the properties cohere and thus support judgments of similarity and generalization across the items that possess them. Conceptual systems must also, however, encode the properties that distinguish birds — the fact that penguins cannot fly, robins have red breasts, parrots are found in the tropics, and so on.

The Complementary Learning Systems (CLS) theory proposes a role for sleep in establishing long-term neocortical representations of such knowledge: The hippocampus rapidly stores new information while the organism is awake, which it then replays during sleep, allowing the slow-learning cortex to gradually incorporate the new information into existing knowledge structures [5]. Specialized subfields of the hippocampus rapidly bind arbitrary new information without interference using sparse, pattern-separated representations [6]. Neocortex, in contrast, employs more overlapping representations, facilitating extraction of the commonalities across individual elements and episodes.

This view suggests that sleep should improve memory for coherent properties: As new learning examples become better integrated in the cortical system, representational overlap should promote cross-generalization amongst conceptually related items, boosting recall of these features. It may seem that idiosyncratic information would be lost in the overlapping cortical representations, and it is of course the case that we tend to forget the details of most of our memories over time. However, hippocampal replay during sleep may provide reminders about these details, helping cortex to represent this information as well, at least relative to a period of time awake.

A substantial literature now suggests that sleep benefits memory for both individual items and relational structure amongst items [7, 8]. Prior research, however, has not assessed the relative fate and prioritization of these memory types during sleep, an understanding of which would provide important constraints on the mechanisms of consolidation. Assessing how sleep differentially impacts these memories requires matching strength of initial encoding, as forgetting rates vary with learning strength [9] and sleep tends to benefit more weakly encoded information [10–17, c.f. 18, 19].

We report two experiments using a novel semantic learning task, in which participants learned both coherent and individuating properties of 15 “satellite” objects organized into three categories. Satellites shared most features with other category members but also possessed unique individuating features. Learning as well as memory assessment for both feature types occurred via property inference [20], a task that captures the semantic system’s primary function of inferring missing properties from partial information [1, 2]. Frequency of exposure during learning was manipulated to ensure that (1) unique and shared properties were acquired equally well, but (2) the strength of this learning prior to sleep varied across the three categories, allowing us to assess how sleep prioritizes semantic memories with differential encoding strength. The paradigm thus employs a single canonical semantic task to assess memory for both coherent and individuating properties, with the degree of initial learning matched across property type but varied across categories.

In Experiment 1, participants in the sleep condition learned at night and were tested before and after sleeping at home overnight, while participants in the wake condition learned in the morning and were tested at the beginning and end of a day awake. In Experiment 2, participants learned and tested in the morning and tested again in the afternoon, either with or without a polysomnographically-recorded nap in between. This allowed us to compare behavioral results of an afternoon nap to overnight sleep and to assess the relationship of different sleep stages to changes in performance.

## Methods

### Participants

In Experiment 1, 111 members of the University of Wisconsin-Madison community (68 females, mean age=19.1 years, range=17–31 years) participated in exchange for course credit or monetary compensation. Data from 9 additional participants were excluded due to performing lower than 2 SD below average on the first session test (2 subjects) and procedural difficulties (7 subjects). The study protocol was approved by the Institutional Review Board at the University of Wisconsin-Madison, and methods were carried out in accordance with all guidelines and regulations.

In Experiment 2, 82 members of the University of California-Riverside community (49 females, mean age=19.9 years, range=18–34 years) participated in exchange for course credit or monetary compensation. Data from 11 additional subjects were excluded due to: napping for less than 20 min (3 subjects), falling asleep during the quiet wake period (7 subjects), and performing lower than 2 SD below average on the first session test (1 subject). Subjects reported having a regular sleep-wake schedule, which was defined as regularly going to bed no later than 2AM, waking up no later than 10AM, and getting at least 7 hours of total sleep per night on average. The Epworth Sleepiness Scale [ESS; 21] and the reduced Morningness-Eveningness Questionnaire [rMEQ; 22] were used to screen out potential subjects with excessive daytime sleepiness (ESS score >10) or extreme chronotypes (rMEQ < 8 or > 21). Experimental procedures were approved by the Human Research Review Board at the University of California at Riverside, and methods were carried out in accordance with all guidelines and regulations. Informed written consent was obtained from all participants in both experiments.

### Stimuli

Participants learned about 15 novel “satellite” objects organized into three classes (Figure 1a). Each satellite had a “class” name (Alpha, Beta, or Gamma) shared with other members of the same category, a unique “code” name (a well-formed nonword), and five visual parts. One of the satellites in each category is the prototype (shown on the left for each category in Figure 1) — it contains all the prototypical parts for that category. Each of the other satellites has one part deviating from the prototype (which part deviates is different for each satellite in the category). Thus, each non-prototype shares 4 features with the prototype and 3 features with other non-prototypes from the same category. Exemplars from different categories do not share any features. Each satellite has *shared features:* the class name and the parts shared among members of the category; it also has *unique features:* the code name and the part unique to that satellite (except for the prototype, which has no unique part). Satellites were constructed randomly for each participant, constrained by this category structure.

**Figure 1.**
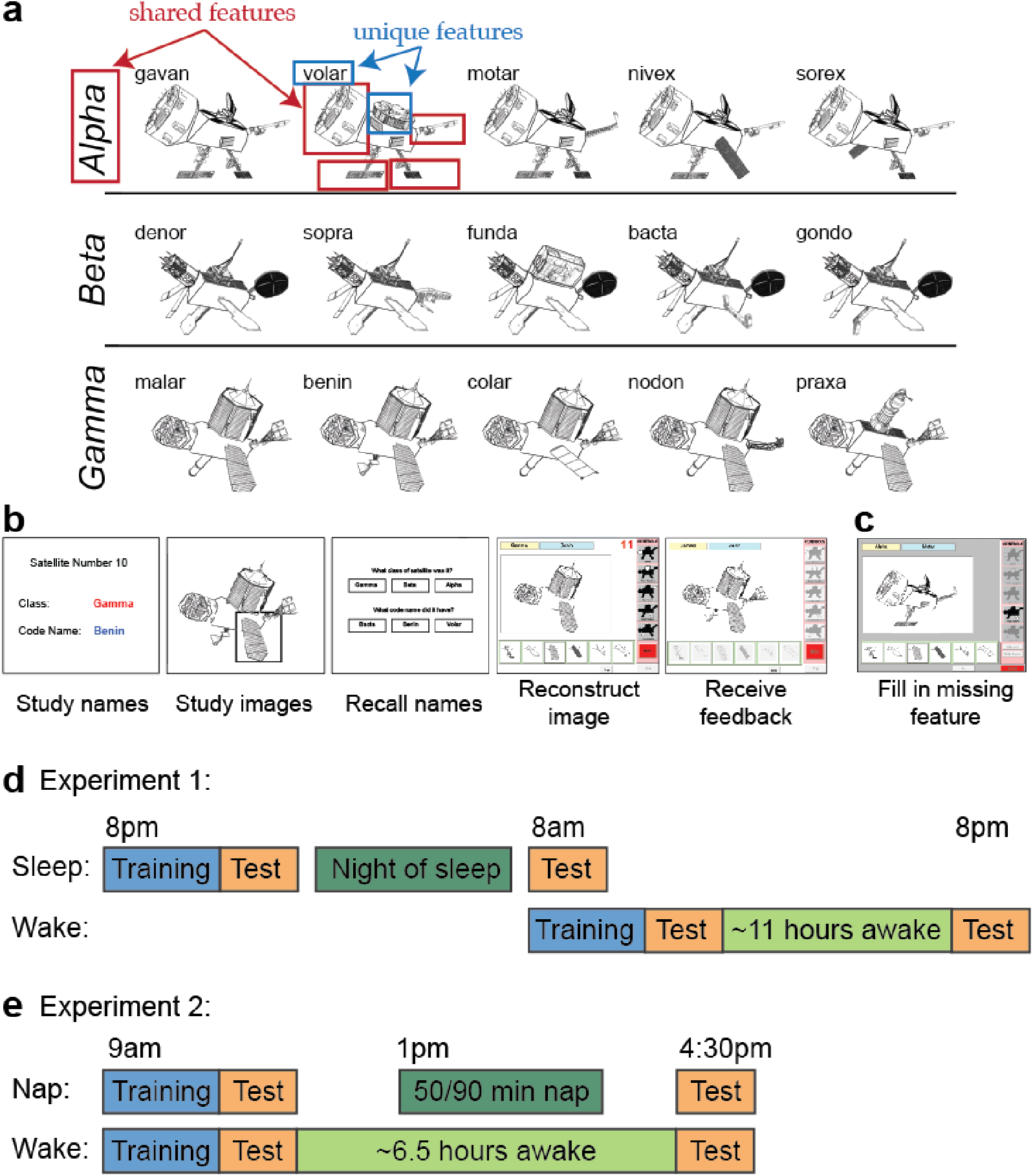
Stimuli and task. (**a**) Examples of satellite stimuli presented from the three classes Alpha, Beta, and Gamma, each labeled with unique code names. Satellites were built randomly for each participant, using the same category structure. Shared and unique features of one satellite are highlighted. (**b**) The one-by-one introduction to each satellite at the beginning of training. (**c**) The second phase of training, in which one feature is missing and participants attempt to fill in the feature, receiving feedback. The test trials are the same but have two features missing. (**d**) Overview of protocol for Experiment 1. (**e**) Overview of protocol for Experiment 2. Time spans in (**d**) and (**e**) are not to scale.

To assess effects of encoding strength, we varied the frequency of exposure of the different categories using a 1-2-3 ratio: 1/6 of all training items were from the low frequency category, 2/6 from the medium frequency category, and 3/6 from the high frequency category.

### Procedure

Participants learned about the satellites in two phases. In the first phase (17 min mean duration), the satellites were introduced one by one (Figure 1b). For each satellite, the class and code name were displayed followed by the satellite image. A box highlighted each visual feature in sequence to encourage participants to attend to each feature. Participants were then asked to recall the class and code names by clicking on one of three options given for each name. Next, participants used a point-and-click interface to try to reconstruct the satellite image from scratch. Icons representing the five part types were displayed on the right hand side of the screen, and when an icon was clicked, all the possible versions of that part were displayed in a row at the bottom of the screen. The participant could then click on one of the part versions on the bottom to add it to the satellite in the center of the screen. If the participant was too slow at this task (took longer than 15 s), or reconstructed the satellite incorrectly, a feedback screen would appear displaying the correct features.

In the second phase (34 min mean duration) participants were shown a satellite with one feature missing (Figure 1c), which could be one of the five visual features, the code name, or the class name (code and class name buttons were displayed along with the part icons on the right hand side of the screen, and when selected, displayed the corresponding name options in a row on the bottom). Using the same point-and-click interface, participants chose a feature amongst six options to complete the satellite. There was no response time deadline. If they chose the correct feature, they were told it was correct, and could move on to the next trial. If they chose an incorrect feature, they were shown the correct feature, and had to repeat the trial until they chose the correct feature.

Remembering the shared properties of the satellites is easier than remembering the unique properties, as the shared properties are reinforced across study of all the satellites in the same class. The task was titrated in pilot testing to ensure that, at the end of training, participants performed equivalently at retrieving shared and unique properties of the satellites. To accomplish this, unique features were queried 24 times more frequently than shared features. This phase of training continued until the participant reached a criterion of 0.66 proportion of trials correct on a block of 32 trials, or until about an hour had passed (cutoff=60 min in Experiment 1, cutoff=75 min in Experiment 2).

Immediately after training, participants were tested by again filling in missing features of the satellites, now without feedback. The test phase had 51 trials, with two missing features per trial, which allowed us to collect more information per trial as well as provide less exposure to the correct features (to minimize learning during the test phase). Each satellite appeared twice in the test phase: once with its code name and its class name or one shared part tested, and once with two shared parts or one unique part and one shared part tested. The remaining 21 trials tested generalization to novel satellites. Novel satellites were members of the trained categories but had one novel feature or a novel combination of features (a prototype from one category with one or two prototypical features swapped in from a different category). The queried feature for novel items was always a shared feature (class name or shared part). Test trials were presented in a random order.

Participants completed the same set of test items (in a different random order) after a delay. They were not told in the first session that there would be a second memory test. The Karolinska Sleepiness Scale [KSS; 23], which assesses state sleepiness/alertness on a scale of 1 (extremely alert) to 9 (very sleepy), was completed at the end of each session.

### Experiment 1-specific procedure

In Experiment 1, 12 hours elapsed between the two sessions (Figure 1d). Participants in the sleep condition (n=61) began the first session around 8pm, and participants in the wake condition (n=50) began the first session around 8am. Participants were given no instructions about their activities between sessions. At the end of the second session, subjects in both groups filled out a questionnaire asking how long they slept between sessions. The KSS was not collected for 22 subjects due to procedural error.

### Experiment 2-specific procedure

Subject participating in this polysomnography (PSG) study underwent stricter sleep screening and procedures. Subjects were instructed to keep a regular sleep schedule and attempt at least 7 hours of sleep per night. Adherence to the sleep schedule was tracked with daily sleep diaries. Additionally, subjects wore an actigraph wrist monitor (Actiwatch Spectrum, Respironics) the night immediately prior to the study day.

Subjects were asked to refrain from consuming caffeine, alcohol, and all stimulants for 24 hours prior to and including the day of the study. Heavy caffeine users (> 240 mg per day) were not enrolled to exclude the possibility of significant withdrawal symptoms during the experiment.

Subjects arrived between 8:30am–9am (Figure 1e). Before proceeding with the experiment, an experimenter checked each subject’s actigraphy data to verify adherence to the sleep schedule the night prior. Session 1 began at approximately 9am.

Upon completion of Session 1, subjects were randomly assigned to one of four groups: active wake (AW), quiet wake (QW), 50 min nap, or 90 min nap. The AW group (n=22) carried out their normal daily activities outside of the lab, but were instructed to abstain from exercise and napping. Wakefulness in the AW group was monitored with the actigraph wrist monitors. Subjects in the QW and nap groups had electrodes attached for standard PSG. At 1pm, the QW group (n=20) commenced listening to short stories on an iPod for 60 min while sitting in a quiet, dark room with PSG monitoring to make sure they did not fall asleep. During QW sessions, an experimenter woke subjects at the first sign of Stage 1 sleep. Subjects in the two nap groups were given a nap opportunity between 1–3PM. If a subject spent more than 30 consecutive minutes awake during the nap window then the nap was ended. Otherwise, an experimenter woke the subject after he or she had obtained the desired amount of total sleep time (50 min or 90 min). Given that shorter naps tend to have less REM sleep than longer naps, the use of these two durations increased the likelihood of having naps with and without REM sleep. Post-hoc sleep stage scoring was used to place subjects into a REM group (n=23, naps contained more than one minute of REM sleep) or non-REM (NREM, n=17) after completion of the experiment. The session 2 test occurred for all groups between 4:30–5pm.

### Polysomnography

PSG data were collected using Astro-Med Grass Heritage Model 15 amplifiers and Grass Gamma software. Eight scalp electroencephalogram (EEG) and two electrooculogram (EOG) electrodes were referenced to unlinked contralateral mastoids (F3/A2, F4/A1, C3/A2, C4/A1, P3/A2, P4/A1, O1/A2, O2/A1, LOC/A2 and ROC/A1), and two electromyogram electrodes were attached under the chin to measure muscle tone. PSG data were digitized at 256 Hz and visually scored in 30 s epochs according to the sleep staging criteria of Rechtschaffen and Kales [24]. Sleep architecture variables included percentage of the nap spent in Stage 1, Stage 2, slow wave sleep (SWS), and rapid eye movement (REM).

EEG data were preprocessed and analyzed using BrainVision Analyzer 2.0 (BrainProducts, Munich Germany) and Matlab. EEG data were bandpass filtered between 0.3 and 35 Hz, and all epochs with artifacts and arousals were identified by visual inspection and rejected. Sleep spindles were automatically detected during Stage 2 and SWS using a wavelet-based algorithm [25]. Following spindle detection, spindle densities were calculated by dividing the number of discrete spindle events by minutes spent in each sleep stage at each scalp EEG electrode site. Data for an individual channel were excluded if the channel was determined to be unreliable.

### Data availability

The data analyzed in this study are included as Supplementary Information.

## Results

### Experiment 1

#### Sleepiness survey

The KSS scores did not differ across sleep and wake conditions for the first session (mean sleep=5.67; mean wake=6.15; *t*[86]=1.406, *p*=0.163) nor second session (mean sleep=5.46; mean wake=4.88, *t*[86]=1.499, *p*=0.138), suggesting that there were no alertness differences between groups due to time of day.

#### Training

Participants trained for an average of 171.55 trials (SD=84.39), including repetition trials for incorrect choices. The average proportion correct on the last training block was 0.686 (SD=0.150).

#### Test performance

For proportion correct on the first test, we found no differences between wake and sleep groups in unique, shared, or novel feature types (*ps*>0.175; Supplementary Fig. S1), indicating no bias due to time of day. We also found no differences between first session unique and shared performance within each of the two groups (*ps*>0.164; Supplementary Fig. S1), demonstrating a successful matching of performance on these feature types. Novel items, which we did not attempt to match in performance, were worse than unique and shared features in the first test: *t*[60]=1.586, *p*=0.118 for the sleep group; *t*[49]=2.987, *p*=0.004 for the wake group.

The frequency manipulation had a robust effect on session 1 performance (Supplementary Fig. S1), which was most pronounced for unique items (LF mean=0.406, SD=0.245; MF mean=0.530, SD=0.235; HF mean=0.659, SD=0.247), followed by shared items (LF mean=0.487, SD=0.229; MF mean=0.535, SD=0.217; HF mean=0.613, SD=0.247), and then novel items (LF mean=0.455, SD=0.208; MF mean=0.472, SD=0.198; HF mean=0.531, SD=0.219). Individual subject slopes across the three frequency levels were significantly greater for unique than shared (*t*[110]=3.983, *p*=0.0001), and marginally greater for shared than novel (*t*[110]=1.687, *p*=0.095).

To assess change in proportion correct from the first to second session, we ran a three way ANOVA on the session 2 – session 1 performances, with sleep vs. wake groups as across-subject factor and frequency and feature type as within-subject factors. There was a main effect of group, with greater (i.e., more positive) change for the sleep group than for the wake group (*F*[1,109]=9.802, *p*=0.002), a main effect of feature type (*F*[2,218]=5.913, *p*=0.003), with unique features improving less than shared and novel, and a main effect of frequency (*F*[2,218]=3.090, *p*=0.047), with lower frequencies faring better (i.e., showing less forgetting) than higher frequencies. Finally, there was a group by feature type interaction (*F*[2,218]=4.074, *p*=0.018), driven by there being no difference between sleep and wake in the novel item features (*t*[109]=0.235, *p*=0.815) but a difference between groups in both unique (*t*[109]=3.098, *p*=0.003) and shared features (*t*[109]=3.267, *p*=0.002). The magnitude of the sleep vs. wake difference for unique vs. shared features was not different, collapsing across frequency (*t*[109]=0.303, *p*=0.763). These results suggest that a night of sleep is much more beneficial than a day awake for memory for unique and shared features of the satellites.

Notably, there was a reliable above-baseline improvement in shared feature memory in the sleep group, collapsing across frequency (Figure 2; *t*[60]=2.093, *p*=0.041; after FDR correction across three feature types, *p*=0.122). This provides evidence that subjects have an improved understanding of the shared category structure after sleeping. Average change in performance for the sleep group was marginally better for shared than unique features (*t*[60]=1.688, *p*=0.097). Breaking the data down by category frequency, the sleep group was not significantly different from zero in any of the nine frequency-specific conditions (*ps*>0.131).

**Figure 2.**
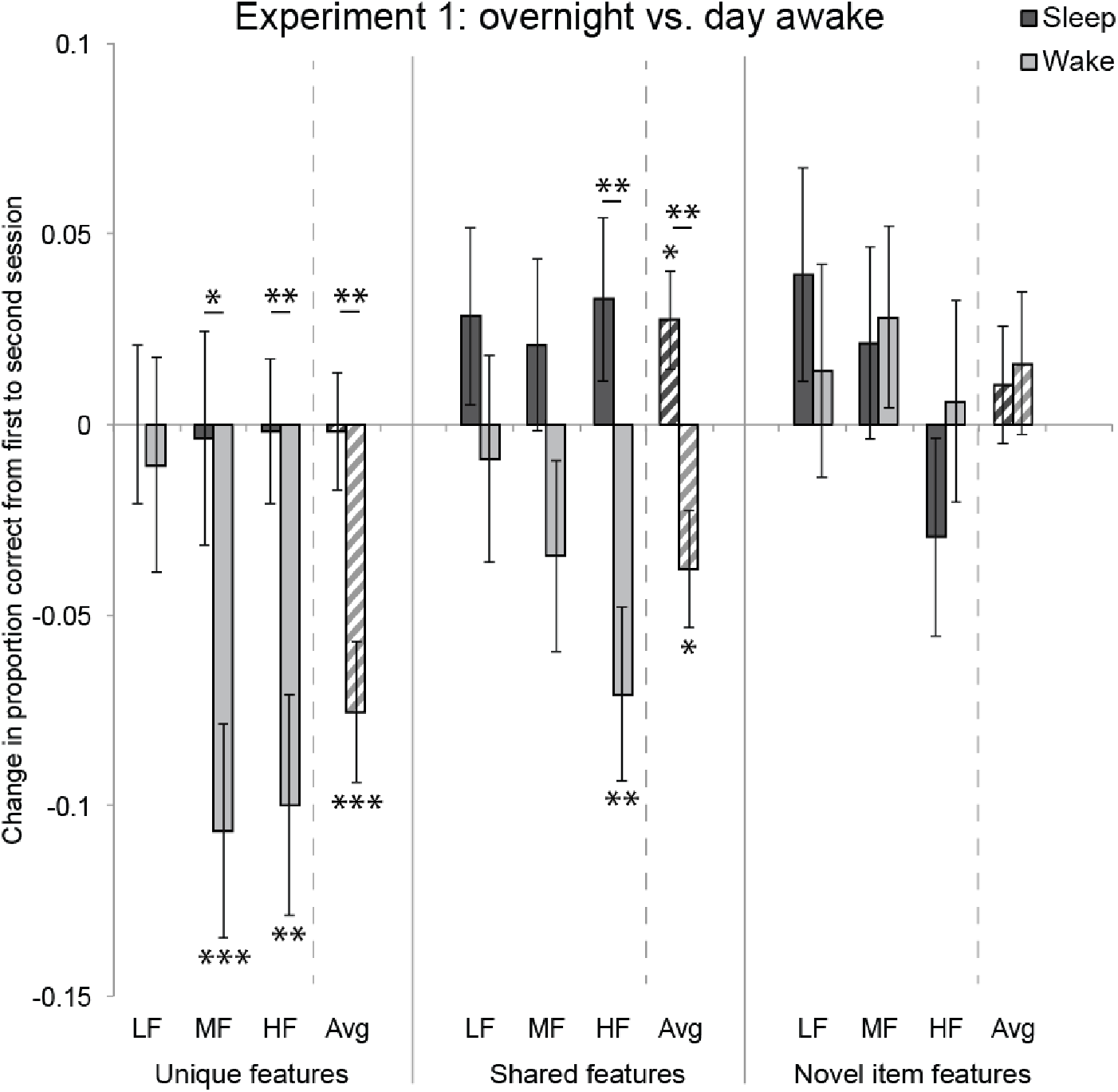
Experiment 1 results. Change in proportion correct from first to second session for unique features, shared features, and novel item features. For each feature type, results are shown for low frequency (LF), medium frequency (MF), and high frequency (HF) category, as well as the average (Avg) across categories. * *p*<0.05, ** *p*<0.01, *** *p*<0.001, uncorrected. Asterisks above horizontal lines show significant differences between conditions; asterisks without bars indicate where conditions differ from zero. Error bars denote ± 1 SEM.

The wake group had reliable forgetting of both unique and shared features, collapsing across frequency (unique: *t*[49]=4.109, *p*=0.0002, FDR-corrected *p*=0.0004; shared: *t*[49]=2.482, *p*=0.017, FDR-corrected *p*=0.026). Breaking the data down by category frequency, there was significant forgetting in MF unique features (*t*[49]=3.805, *p*=0.0004, FDR-corrected *p*=0.004), HF unique features (*t*[49]=3.452, *p*=0.001, FDR-corrected *p*=0.005), and HF shared features (*t*[49]=3.105, *p*=0.003, FDR-corrected *p*=0.010). No other conditions differed significantly from zero (*ps*>0.173). Forgetting was reliably greater in MF and HF unique features than in LF unique features (MF: *t*[49]=2.117, *p*=0.039; HF: *t*[49]=2.017, *p*=0.049). Forgetting was also reliably greater in HF shared features than LF shared features (*t*[49]=2.038, *p*=0.047).

#### Verbal vs. visual information

Each feature type had verbal and visual subtypes: Unique features could be a visual part or a verbal code name, shared features could be a visual part or a verbal class name, and novel item features could be a visual part or a verbal class name. To assess whether verbal vs. visual type had any effect, we re-ran the ANOVA with verbal vs. visual as an additional within-subject factor and found a large main effect of this factor (*F*[1,109]=18.794, *p*=0.00003), with verbal information remembered better than visual across the delay. There were no interactions between this factor and group (or any other factors), however, suggesting that sleep does not have a different effect depending on whether the feature is learned verbally vs. visually.

#### Nap survey

Out of 50 subjects in the wake group, 15 reported taking a nap between the first and second session. The mean nap length was 64.0 minutes (median=60, SD=35.36, range=15–135). These subjects did not differ from those who did not nap in their change from first to second session performance for any feature type at any frequency level (*ps*>0.138), suggesting that any benefit of the nap was swamped by the effect of a much longer period of time awake. Including subjects who napped in the wake group might have been expected to reduce the magnitude of the effects, but Cohen’s *d* values were slightly stronger (more negative) for unique (nap *d*=-0.564, no nap *d*=-0.356) and shared features (nap *d*=-0.310, no nap *d*=-0.182) in the group that napped, collapsing across frequency (for novel item features, nap *d*=0.140, no nap *d*=0.075). We therefore collapsed across these two subgroups for all reported analyses.

#### Discussion

The overall picture from Experiment 1 is that a night of sleep promotes retention of the unique features of category members and improves memory for shared features, whereas both of these feature types are forgotten across a day awake. In the wake group, the forgetting effect was larger for higher-frequency items, which may in part reflect a proportional forgetting function (i.e., the better memory is initially, the larger forgetting will be). The sleep group resisted this forgetting effect, across all frequency levels. There were no changes or group effects in generalization to novel item features (see General Discussion for possible explanations).

We next ran a nap variant of the paradigm with polysomnography, allowing us to explore the potential influence of sleep features and the effects of a shorter sleep/wake period, with tests at matched times of day across groups.

### Experiment 2

#### Sleepiness survey

In the first session, there was no difference in KSS scores between the groups that did sleep versus those that did not (mean sleep groups=4.025; mean wake groups=4.357; *t*[80]=0.803, *p*=0.425). In the second session, the groups that slept reported being less sleepy than the groups that did not (mean sleep groups=2.700; mean wake groups=4.381; *t*[80]=4.524, *p*<0.001). However, KSS scores in the second session did not correlate, in either group, with second session performance or change from first to second session performance for any feature type (*ps*>0.123).

#### Training

Participants trained for an average of 199.96 trials (SD=108.23). Average performance on the last training block was 0.748 (SD=0.060).

#### Test performance

For proportion correct on the first test, we found no differences between quiet wake, active wake, NREM, or REM groups in unique, shared, or novel feature types (*ps*>0.150), indicating no bias across groups before the delay. We also found no differences between first session unique and shared performance within any of the four groups (*ps*>0.172). Novel items, which we did not attempt to match in performance, were worse than unique and shared features in the first test: *t*[16]=4.447, *p*<0.001 for NREM; *t*[22]=4.803, *p*<0.001 for REM; *t*[21]=3.095, *p*=0.006 for AW; *t*[19]=1.829, *p*=0.083 for QW.

Looking at the change in proportion correct from the first to second session, we found no differences between QW and AW, nor between NREM and REM groups, for any frequency level or feature type (*ps*>0.076). We therefore collapsed the groups into nap (NREM and REM) and wake (QW and AW) for further analyses, except when assessing the contribution of sleep stages.

The frequency manipulation had a robust effect on session 1 performance (Supplementary Fig. S2), which was again most pronounced for unique items (LF mean=0.391, SD=0.200; MF mean=0.577, SD=0.199; HF mean=0.745, SD=0.189), followed by shared items (LF mean=0.537, SD=0.198; MF mean=0.603, SD=0.202; HF mean=0.646, SD=0.207), and then novel items (LF mean=0.477, SD=0.180; MF mean=0.460, SD=0.192; HF mean=0.494, SD=0.209). Individual subject slopes across the three frequency levels were significantly greater for unique than shared (*t*[81]=6.690, *p*<0.0001), and significantly greater for shared than novel (*t*[81]=2.629, *p*=0.010).

The sleep and wake nap groups did not differ from the sleep and wake groups in Experiment 1 in first session performance, for any frequency or item type (*ps*>0.0667), but the pattern of change over time was different. We again ran a three way ANOVA on session 2 – session 1 performance, with nap vs. wake groups as across-subject factor and frequency and feature type as within-subject factors. There was a main effect of feature type (*F*[2,160]=10.450, *p*=0.00005), a feature type by frequency interaction (*F*[4,320]=3.104, *p*=0.016), and, critically, a feature type by frequency by group interaction (*F*[4,320]=3.104, *p*=0.016). This indicates that there was no overall effect of nap versus wake, nor differences between the groups for unique, shared, and novel feature types, but rather that the groups differed in certain frequencies and feature types. In particular, the nap group improved more than the sleep group on LF shared features (*t*[80]=2.846, *p*=0.006; with FDR correction across 3 feature types, 3 frequencies, *p*=0.051; Figure 3). This may reflect a prioritization of the category that is least well learned.

**Figure 3.**
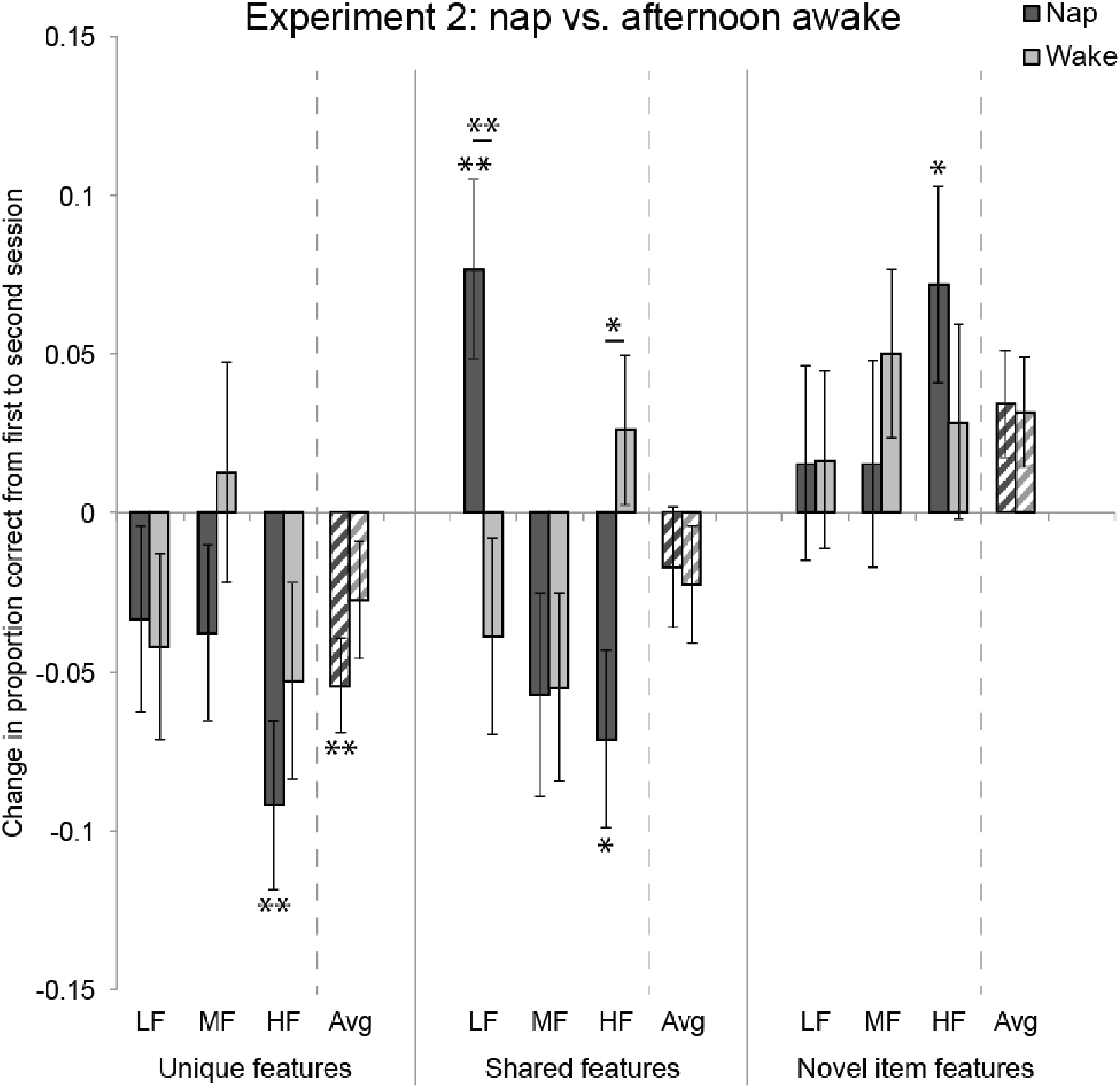
Experiment 2 results. Same structure as in Figure 2. * *p*<0.05, ** *p*<0.01, uncorrected.

Notably, as in Experiment 1, this change in LF shared feature performance reflected an above-baseline improvement for the nap group — the sleep group performed better on LF shared features after the nap than before (*t*[39]=2.888, *p*=0.006). There was also a difference between the nap and wake groups for high frequency shared features (*t*[80]=2.583, *p*=0.012; with FDR correction, *p*=0.052), with wake better than sleep, though this did not reflect above baseline improvement for the wake group (*t*[41]=1.105, *p*=0.275). The feature type by frequency interaction reflects the fact that there was more forgetting, across both groups, for higher frequency unique and shared features (but not for novel item features).

#### Verbal vs. visual information

We again tested for effects of verbal vs. visual feature type by re-running the ANOVA with verbal vs. visual as an additional within subject factor. We found a main effect (*F*(1,80)=4.469, *p*=0.038), with verbal information again remembered better than visual across the delay. There were again no interactions between this factor and group (or any other factors).

#### Sleep features

To assess the potential contribution of different sleep stages to the improvement in shared features in the low frequency category, we first looked at performance separately for the NREM and REM groups (Figure 4a). The NREM group improved significantly from session 1 to 2 (*t*[16]=2.954, *p*=0.009), suggesting that NREM sleep alone is sufficient for this change in performance. The REM group was no different than the NREM group (*t*[38]=0.525, *p*=0.603), but was not itself reliably above baseline (*t*[22]=1.602, *p*=0.123).

**Figure 4.**
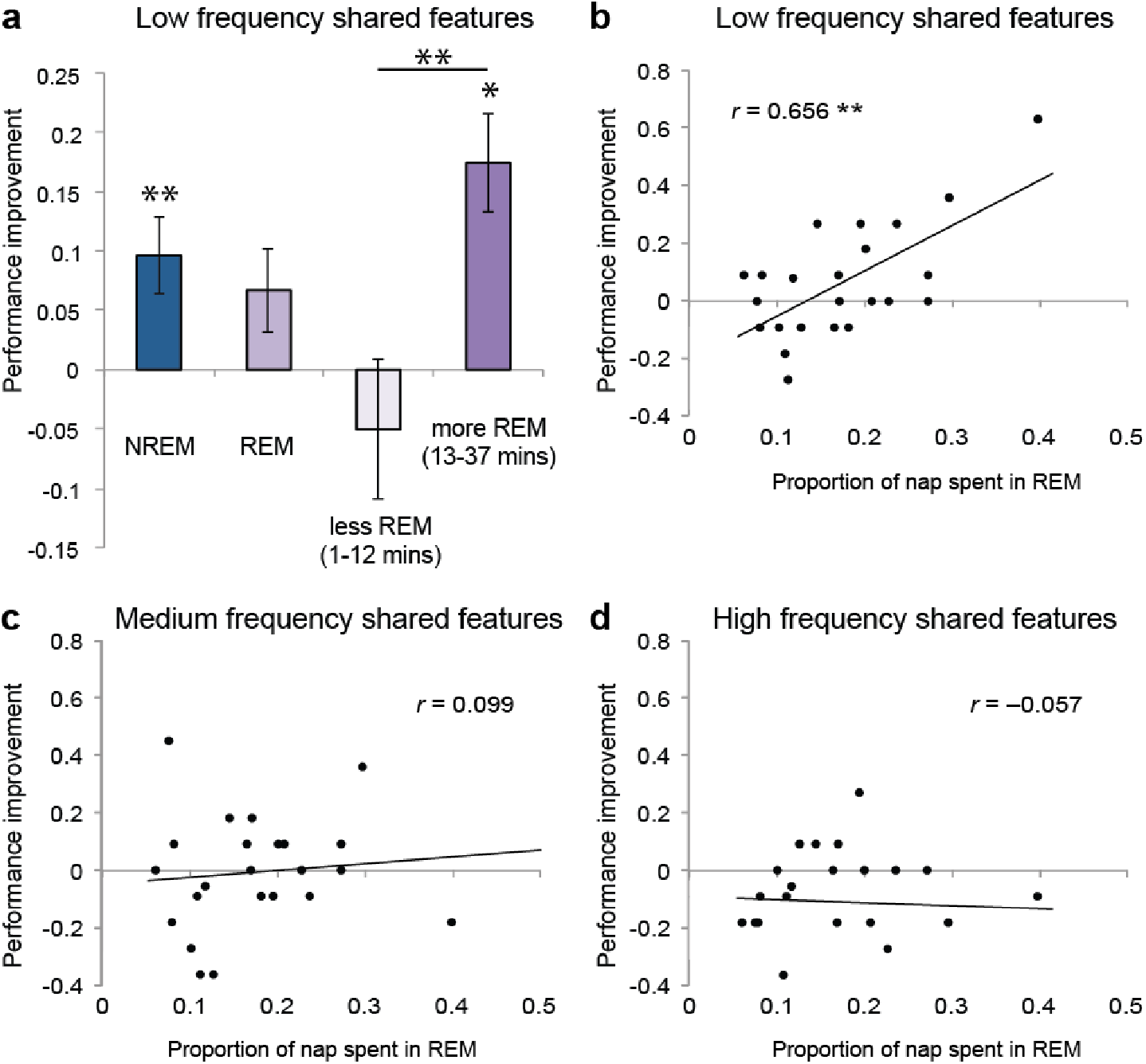
Relationship between NREM and REM sleep and behavioral change. (**a**) Improvement in low frequency shared features for the NREM group, the REM group, and for REM groups on either side of a median split by number of minutes in REM. (**b, c, d**) Relationship between proportion of nap spent in REM and performance improvement for low frequency, medium frequency, and high frequency shared features. * *p*<0.05, ** *p*<0.01.

We next assessed whether amount of REM sleep is related to change in performance. We found that among the subjects who had REM sleep there was a strong correlation between proportion of time in the nap spent in REM sleep and performance improvement (Figure 4b; *r*=0.656, *p*=0.001). This was not true for shared features in medium or high frequency categories (Figure 4c,d); The correlation between REM and performance improvement was reliably greater for LF than MF (*p*=0.03) and HF (*p*=0.008) shared features, suggesting that the nap indeed focused on processing of shared features in the low frequency category. Improvement in low frequency shared features was not correlated with total sleep time (*r*=−0.002, *p*=0.989), suggesting that this is not simply about sleeping longer, but about time spent in REM.

We also performed a median split analysis on number of minutes spent in REM sleep, which divided the group into subjects getting 1–12 minutes and those getting 13–37 minutes (Figure 1a). Subjects who get less REM showed a numerical decrement in performance, whereas those with more REM performed numerically better than the NREM group (difference between less REM and more REM: *t*[21]=3.212, *p*=0.004). This pattern suggests that a small amount of REM is actually worse than having no REM at all (less REM vs. NREM: *t*[26]=2.966, *p*=0.006).

We also performed exploratory analyses for a collapsed NREM and REM group, correlating unique, shared, and novel feature improvement at each of the three frequency levels, for stage 1, stage 2, SWS, and total sleep time. We did not find any reliable correlations (all uncorrected *ps*>0.073). We performed further exploratory analyses assessing whether spindle density in stage 2 or SWS at any channel correlated with improvement for any of the three feature types, at any of the three frequency levels. While there were reliable uncorrected correlations, none survived FDR correction (*ps*>0.138).

#### Discussion

Over a short period of sleep, we again found an above-baseline improvement in shared feature memory, but only for the LF category — the category that starts off with the poorest performance. This suggests a potential prioritization of the weakest category early in a sleep period.

There were no differences between NREM and REM groups in terms of behavioral change, nor between active wake and quiet wake, in contrast to prior studies [e.g., 14]. When considering the PSG data, the group with only NREM sleep showed reliable improvement in LF shared features, suggesting that NREM sleep is sufficient to promote better category understanding, and REM sleep time was associated with the improvement, suggesting that REM is also doing relevant processing.

As in Experiment 1, memory declined more for the higher frequency categories, which start off better learned. Unlike Experiment 1, the frequency-influenced forgetting occurred for both sleep and wake groups (with the puzzling exception of HF shared features in the wake group). We again found no notable changes in novel item features.

## General Discussion

Effective consolidation of new semantic learning must simultaneously encompass both object-specific information and the conceptual structure existing across objects. Experiment 1 showed that a night of sleep indeed benefits both knowledge types: Whereas shared and unique features showed forgetting (especially for well-learned items) over wake, memory was preserved for unique features and improved for shared features over sleep regardless of initial encoding strength. Experiment 2 found memory improvements for lower-frequency shared features in a nap paradigm, with polysomnography suggesting that both NREM and REM phases contributed to these gains. In this section we consider the implications of these results for our understanding of semantic knowledge consolidation.

### Enhancement versus maintenance of new learning

The existing sleep literature has consistently suggested that, though sleep may enhance non-declarative learning, it only serves to prevent or decrease forgetting of declarative memory, perhaps just passively protecting against interference [26]. Contrary to this view, we observed above-baseline improvement in memory for shared features in a declarative memory task, suggesting an active consolidation process [27]. What accounts for the difference? Prior studies of declarative memory in the sleep literature have mainly used episodic memory tasks, requiring arbitrary associations (e.g., word pairs or object locations). Our results suggest that the division between what is maintained versus enhanced by sleep may not be about declarative vs. nondeclarative information, but instead about arbitrary versus structured information. Our unique feature condition, in which we found a prevention of forgetting overnight, is more episodic-like, in the sense that the associations are arbitrary, and may correspond to the kinds of processing observed in prior episodic memory sleep studies. Our shared feature condition, like prior studies of non-declarative memory, would be expected to benefit from a stronger cortical representation. Furthermore, other studies of structured information have also found above-baseline improvement [16, 28–31]. The current data suggest that, for new semantic learning, sleep promotes maintenance of arbitrary information and simultaneous enhancement of memory for representational structure.

### Prioritization of weak memories

In Experiment 2, which had shorter sleep and wake intervals between tests, we again saw a true improvement in shared feature memory in the sleep group, but only for the category that had been exposed at the lowest frequency during training, with forgetting occurring for the high frequency category. This finding is highly consistent with an accumulating literature suggesting that weaker memories are prioritized during sleep [10–17], although some studies have found a focus on stronger memories [18, 19]. Combined with Experiment 1, the results suggest that sleep first prioritizes the information most in need of help, and then moves on to other information. In fact, in Experiment 1, the largest difference between sleep and wake groups was in the high frequency category, suggesting a potential prioritization of stronger information later in the night. The mechanism by which weaker memories are tagged for early prioritization is an important topic for future research. Subjects in these studies did not receive feedback during the test phase, so tagging must have relied on some internal sense of performance accuracy.

Experiment 2 also suggests that initial strength of encoding is not the only factor determining the subsequent fate of memories over sleep or wake. Low frequency unique features did not benefit from a nap despite being more weakly encoded than low frequency shared features. That is, sleep only benefited weakly-learned properties shared amongst category members. Thus, the degree to which sleep appears to prioritize some memories over others depends not only on strength of initial learning and amount of sleep, but, again, on the degree to which the information is structured.

### Forgetting of unique features differs for nap versus full night sleep

Experiments 1 and 2 both found that memory for unique features declined over wake, more so in Experiment 1 where the delay between initial and final test was longer. In both studies, low frequency unique features were preserved over sleep. The pattern was very different, however, for high frequency unique features: A full night’s sleep produced good retention of unique features regardless of frequency, while a short nap produced significant forgetting of high frequency unique features. The pattern makes sense if early sleep prioritizes weaker memories — without active processing during the nap, the stronger items are forgotten as if the participant had stayed awake, but more sleep (or more NREM-REM cycles) provides opportunities for rescue. The combined results thus provide evidence that unique, in addition to shared features, are processed in an active way during sleep. In general, the differences between the nap and overnight studies suggest that for this paradigm, unlike others [e.g., 32], a nap is not equivalent to a full night of sleep — a shorter sleep opportunity in a rich object category learning paradigm may only allow focus on items most in need of help.

### Active gains occur in both NREM and REM sleep

Experiment 2 found that NREM sleep alone improves memory for low frequency shared features. The REM group showed numerically smaller benefits, but the amount of REM sleep was strongly positively correlated with improvement — a relationship specific to low frequency shared features. The data suggest that NREM plus sufficient REM sleep most strongly benefits knowledge of category structure, but that NREM plus a small amount of REM can degrade such knowledge. Both NREM and REM sleep have previously been associated with memory for structured information. The amount of NREM sleep correlates with associative inference [33] and statistical learning of tone sequences [28], and the amount of REM sleep is associated with improved performance on the remote associates task [34]. Both sleep stages have been associated with learning a hidden linguistic rule [35] and probabilistic learning on the weather prediction task [16, 36; note that this paper suggests that the association is based on trait-based REM duration]. NREM sleep may be the time when the hippocampus replays new information to cortex, while REM helps to further stabilize the new information in cortical networks [37]. Both of these processes would be expected to benefit understanding of shared structure. The possibility that conceptual structure benefits most from hippocampal replay followed by cortical stabilization is a hypothesis that we hope to address in future work.

### Sleep does not influence generalization in the property inference task

In neither experiment did we observe differences between sleep and wake groups in generalization to novel satellites. This was surprising, as a better understanding of shared category structure would be expected to lead to better ability to generalize [2], and prior studies have found benefits of sleep on generalization [29–31, 38–42]. It is possible that a longer period of time would need to elapse before seeing generalization in this paradigm [see 43], or that our assessment was not sensitive enough: Generalization was better than chance but reliably worse than memory for unique and shared properties. It is also possible that there was variance across participants in the strategies used for making judgments about never-before-seen features that hindered our ability to detect generalization.

If the lack of a sleep benefit for generalization is not due to insensitivity or variance in strategy use, how can we interpret this null result? One possibility is that the improvement in memory for shared features in the sleep group does not reflect cortical learning of category structure, but instead improved hippocampal memory for individual satellites. Shared feature memory can be supported by hippocampal episodic memory (in addition to cortical structure learning), and thus improved memory for individual satellites could be supported by strengthened hippocampal item traces without new cortical learning. Under this account, shared features exhibit more robust improvement than unique features because cortical category structure (acquired during initial wake learning, but not boosted by sleep) would serve to selectively amplify hippocampal recall of shared (vs. unique) features. Adjudicating between this account (hippocampal strengthening without cortical learning during sleep) and accounts that posit cortical structure learning during sleep will require further work carefully assessing generalization in this paradigm.

### Other related work

There have been a few prior studies finding effects of sleep on semantic memory [41, 44], but they have focused on integrating new information with existing semantic memory networks, not learning an entirely novel domain, as in our study. One study that did look at novel conceptual learning found retention of memory for category exemplars as well as retention of ability to generalize to novel exemplars and never-seen prototypes across a night of sleep, but not a day awake, in a dot pattern categorization task [45]. This study is consistent with ours in suggesting a benefit of overnight sleep for both unique and shared structure. They did not find any reliable above-baseline improvements, but there was numerical improvement in novel exemplar accuracy similar in magnitude to our shared feature effects, and statistical power may have been lower due to an across-subject design and fewer subjects.

Another study found increased generalization in an object categorization task over the course of an afternoon delay in both nap and wake conditions, with no difference between the two [46]. The task assessed memory or inference for the locations of faces, where some locations were predicted by feature rules (e.g. faces at one location were all young, stout, and had no headwear) and other locations had no rules. Overall memory for studied faces decreased over time, but more so for faces at locations without feature rules, suggesting a benefit due to shared structure in the rule location. While there are many differences between this paradigm and ours, our findings suggest the possibility that the forgetting and lack of sleep-wake differences observed may be due to averaging across all memories, instead of focusing on the weaker memories; Our Experiment 2 findings averaged across frequency are qualitatively similar to the findings from this study.

### How does semantic memory consolidation work in the brain?

The current results are consistent with the idea that initial memories of object categories are hippocampally dependent but that sleep serves to increase reliance on cortical representations. As cortical representations are more overlapping, they are better suited to representing shared structure, allowing an above-baseline benefit not seen for object-unique features. Because our task involves learning across visual and verbal modalities, we would expect the anterior temporal lobe to be the site of such cortical consolidation [47]. Another possibility to consider is that consolidation occurring within the hippocampus itself is responsible for the improvement in shared feature memory. There is certainly hippocampal plasticity during sleep [e.g., 48], and the CA1 subfield of the hippocampus, implicated in consolidation [49], may be capable of representing coherent structure using a moderately overlapping neural code [50]. Lastly, as noted above, the lack of sleep benefits on generalization leaves open another possibility, whereby shared feature memory is improved through strengthened hippocampal item traces interacting with overlapping representations stored (but not locally strengthened) in cortex. Future research will be needed to disentangle these possibilities.

Our findings also contribute to our understanding of semantic memory consolidation mechanisms in suggesting that sleep first prioritizes objects that are weakly learned and most in need of further processing, preventing forgetting of unique features and promoting knowledge of shared features for such items. Longer sleep leads to maintenance of unique features and enhancement of shared category features regardless of initial strength of learning. Overall, our results suggest that sleep actively shapes learning of semantic category structure while simultaneously preserving knowledge of individuating details.

## Author contributions

ACS, KAN, and TTR designed the study. LC managed data collection for Experiment 1. EAM managed data collection and analyzed polysomnography data for Experiment 2. ACS and TTR analyzed behavioral data from both experiments. All authors discussed the interpretation of the data. ACS drafted the manuscript and all authors edited the manuscript.

## Competing financial interests

The authors declare no competing financial interests.

## Acknowledgments

We thank Roy Cox and Robert Stickgold for helpful discussions and Christopher Cox, Nicholas Reihanabad, and Chalani Perera for help running subjects. This work was supported by: NIH NINDS F32-NS093901 (ACS); NSF GRFP (EAM); NIH NIA R01-AG046646 (SCM); NSF BCS-1439210 (SCM); NIH NIMH R01-MH069456 (KAN).

## References

1. Tenenbaum, J.B., Griffiths, T.L., and Kemp, C. (2006). Theory-based Bayesian models of inductive learning and reasoning. Trends Cogn Sci 10, 309–318.

2. Rogers, T.T., and McClelland, J.L. (2004). Semantic Cognition: A Parallel Distributed Processing Approach. (Cambridge, MA: MIT Press).

3. Keil, F.C. (1989). The MIT Press series in learning, development, and conceptual change. Concepts, kinds, and cognitive development. (Cambridge, MA: The MIT Press).

4. Murphy, G.L., and Medin, D.L. (1985). The role of theories in conceptual coherence. Psychol Rev 92, 289–316.

5. McClelland, J.L., McNaughton, B.L., and O’Reilly, R.C. (1995). Why there are complementary learning systems in the hippocampus and neocortex: insights from the successes and failures of connectionist models of learning and memory. Psychol Rev 102, 419–457.

6. Norman, K.A., and O’Reilly, R.C. (2003). Modeling hippocampal and neocortical contributions to recognition memory: a complementary-learning-systems approach. Psychol Rev 110, 611–646.

7. Rasch, B., and Born, J. (2013). About sleep’s role in memory. Physiological reviews 93, 681–766.

8. Landmann, N., Kuhn, M., Piosczyk, H., Feige, B., Baglioni, C., Spiegelhalder, K., Frase, L., Riemann, D., Sterr, A., and Nissen, C. (2014). The reorganisation of memory during sleep. Sleep medicine reviews 18, 531–541.

9. Loftus, G.R. (1985). Evaluating forgetting curves. Journal of Experimental Psychology: Learning, Memory, and Cognition 11, 397–406.

10. Drosopoulos, S., Schulze, C., Fischer, S., and Born, J. (2007). Sleep’s function in the spontaneous recovery and consolidation of memories. J Exp Psychol Gen 136, 169–183.

11. Peters, K.R., Smith, V., and Smith, C.T. (2007). Changes in sleep architecture following motor learning depend on initial skill level. J Cogn Neurosci 19, 817–829.

12. Diekelmann, S., Born, J., and Wagner, U. (2010). Sleep enhances false memories depending on general memory performance. Behav Brain Res 208, 425–429.

13. Cairney, S.A., Lindsay, S., Sobczak, J.M., Paller, K.A., and Gaskell, M.G. (2016). The Benefits of Targeted Memory Reactivation for Consolidation in Sleep are Contingent on Memory Accuracy and Direct Cue-Memory Associations. Sleep 39, 1139–1150.

14. McDevitt, E.A., Duggan, K.A., and Mednick, S.C. (2015). REM sleep rescues learning from interference. Neurobiology of learning and memory 122, 51–62.

15. Kuriyama, K., Stickgold, R., and Walker, M.P. (2004). Sleep-dependent learning and motor-skill complexity. Learning & memory 11, 705–713.

16. Djonlagic, I., Rosenfeld, A., Shohamy, D., Myers, C., Gluck, M., and Stickgold, R. (2009). Sleep enhances category learning. Learning & memory 16, 751–755.

17. Sio, U.N., Monaghan, P., and Ormerod, T. (2013). Sleep on it, but only if it is difficult: effects of sleep on problem solving. Mem Cognit 41, 159–166.

18. Tucker, M.A., and Fishbein, W. (2008). Enhancement of declarative memory performance following a daytime nap is contingent on strength of initial task acquisition. Sleep 31, 197–203.

19. Talamini, L.M., Nieuwenhuis, I.L., Takashima, A., and Jensen, O. (2008). Sleep directly following learning benefits consolidation of spatial associative memory. Learning & memory 15, 233–237.

20. Chin-Parker, S., and Ross, B.H. (2002). The effect of category learning on sensitivity to within-category correlations. Mem Cognit 30, 353–362.

21. Johns, M.W. (1992). Reliability and factor analysis of the Epworth Sleepiness Scale. Sleep 15, 376–381.

22. Adan, A., and Almirall, H. (1991). Horne & Ostberg morningness-eveningness questionnaire: a reduced scale. Personality and Individual Differences 12, 241–253.

23. Akerstedt, T., and Gillberg, M. (1990). Subjective and objective sleepiness in the active individual. Int J Neurosci 52, 29–37.

24. Rechtschaffen, A., and Kales, A. (1968). A Manual of Standardized Terminology, Techniques, and Scoring Systems for Sleep Stages of Human Subjects. (Los Angeles: Brain Information/Brain Research Institute UCLA).

25. Wamsley, E.J., Tucker, M.A., Shinn, A.K., Ono, K.E., McKinley, S.K., Ely, A.V., Goff, D.C., Stickgold, R., and Manoach, D.S. (2012). Reduced sleep spindles and spindle coherence in schizophrenia: mechanisms of impaired memory consolidation? Biol Psychiatry 71, 154–161.

26. Mednick, S.C., Cai, D.J., Shuman, T., Anagnostaras, S., and Wixted, J.T. (2011). An opportunistic theory of cellular and systems consolidation. Trends Neurosci 34, 504–514.

27. Ellenbogen, J.M., Payne, J.D., and Stickgold, R. (2006). The role of sleep in declarative memory consolidation: passive, permissive, active or none? Current opinion in neurobiology 16, 716–722.

28. Durrant, S.J., Taylor, C., Cairney, S., and Lewis, P.A. (2011). Sleep-dependent consolidation of statistical learning. Neuropsychologia 49, 1322–1331.

29. Ellenbogen, J.M., Hu, P.T., Payne, J.D., Titone, D., and Walker, M.P. (2007). Human relational memory requires time and sleep. Proc Natl Acad Sci U S A 104, 7723–7728.

30. Frost, R.L., and Monaghan, P. (2017). Sleep-Driven Computations in Speech Processing. PloS one 12, e0169538.

31. Fenn, K.M., Nusbaum, H.C., and Margoliash, D. (2003). Consolidation during sleep of perceptual learning of spoken language. Nature 425, 614–616.

32. Mednick, S., Nakayama, K., and Stickgold, R. (2003). Sleep-dependent learning: a nap is as good as a night. Nat Neurosci 6, 697–698.

33. Lau, H., Tucker, M.A., and Fishbein, W. (2010). Daytime napping: Effects on human direct associative and relational memory. Neurobiology of learning and memory 93, 554–560.

34. Cai, D.J., Mednick, S.A., Harrison, E.M., Kanady, J.C., and Mednick, S.C. (2009). REM, not incubation, improves creativity by priming associative networks. Proc Natl Acad Sci U S A 106, 10130–10134.

35. Batterink, L.J., Oudiette, D., Reber, P.J., and Paller, K.A. (2014). Sleep facilitates learning a new linguistic rule. Neuropsychologia 65, 169–179.

36. Lerner, I., Lupkin, S.M., Corter, J.E., Peters, S.E., Cannella, L.A., and Gluck, M.A. (2016). The influence of sleep on emotional and cognitive processing is primarily trait-(but not state-) dependent. Neurobiology of learning and memory 134 Pt B, 275–286.

37. Diekelmann, S., and Born, J. (2010). The memory function of sleep. Nat Rev Neurosci 11, 114–126.

38. Nieuwenhuis, I.L., Folia, V., Forkstam, C., Jensen, O., and Petersson, K.M. (2013). Sleep promotes the extraction of grammatical rules. PloS one 8, e65046.

39. Gomez, R.L., Bootzin, R.R., and Nadel, L. (2006). Naps promote abstraction in language-learning infants. Psychol Sci 17, 670–674.

40. Tamminen, J., Davis, M.H., Merkx, M., and Rastle, K. (2012). The role of memory consolidation in generalisation of new linguistic information. Cognition 125, 107–112.

41. Lau, H., Alger, S.E., and Fishbein, W. (2011). Relational memory: a daytime nap facilitates the abstraction of general concepts. PloS one 6, e27139.

42. Pace-Schott, E.F., Milad, M.R., Orr, S.P., Rauch, S.L., Stickgold, R., and Pitman, R.K. (2009). Sleep promotes generalization of extinction of conditioned fear. Sleep 32, 19–26.

43. Lutz, N.D., Diekelmann, S., Hinse-Stern, P., Born, J., and Rauss, K. (2017). Sleep Supports the Slow Abstraction of Gist from Visual Perceptual Memories. Scientific reports 7, 42950.

44. Tamminen, J., Lambon Ralph, M.A., and Lewis, P.A. (2013). The role of sleep spindles and slow-wave activity in integrating new information in semantic memory. J Neurosci 33, 15376–15381.

45. Graveline, Y.M., and Wamsley, E.J. (2017). The impact of sleep on novel concept learning. Neurobiology of learning and memory 141, 19–26.

46. Sweegers, C.C., and Talamini, L.M. (2014). Generalization from episodic memories across time: a route for semantic knowledge acquisition. Cortex 59, 49–61.

47. Lambon Ralph, M.A., Jefferies, E., Patterson, K., and Rogers, T.T. (2017). The neural and computational bases of semantic cognition. Nat Rev Neurosci 18, 42–55.

48. Grosmark, A.D., Mizuseki, K., Pastalkova, E., Diba, K., and Buzsaki, G. (2012). REM sleep reorganizes hippocampal excitability. Neuron 75, 1001–1007.

49. Remondes, M., and Schuman, E.M. (2004). Role for a cortical input to hippocampal area CA1 in the consolidation of a long-term memory.Nature 431, 699–703.

50. Schapiro, A.C., Turk-Browne, N.B., Botvinick, M.M., and Norman, K.A. (2017). Complementary learning systems within the hippocampus: a neural network modelling approach to reconciling episodic memory with statistical learning. Philosophical transactions of the Royal Society of London. Series B, Biological sciences 372.

